# The current landscape and emerging challenges of benchmarking single-cell methods

**DOI:** 10.1101/2023.12.19.572303

**Authors:** Yue Cao, Lijia Yu, Marni Torkel, Sanghyun Kim, Yingxin Lin, Pengyi Yang, Terence P Speed, Shila Ghazanfar, Jean Yee Hwa Yang

## Abstract

With the rapid development of computational methods for single-cell sequencing data, benchmarking serves as a valuable resource. As the number of benchmarking studies surges, it is timely to assess the current state of the field. We conducted a systematic literature search and assessed 282 papers, including all 130 benchmark-only papers from the search and an additional 152 method development papers containing benchmarking. This collective effort provides the most comprehensive quantitative summary of the current landscape of single-cell benchmarking studies. We examine performances across nine broad categories, including often ignored aspects such as role of datasets, robustness of methods and downstream evaluation. Our analysis highlights challenges such as how to effectively combine knowledge across multiple benchmarking studies and in what ways can the community recognise the risk and prevent benchmarking fatigue. This paper highlights the importance of adopting a community-led research paradigm to tackle these challenges and establish best practice standards.

## Introduction

Single-cell sequencing has gained tremendous popularity in recent years and there has been an explosion of computational methods for analysing single-cell data since 2017. Within the domain of single-cell RNA-sequencing (scRNA-seq) alone, there are now over 1500 tools that have been recorded at www.scrna-tools.org (Zappia and Theis 2021). The exponential growth in single-cell RNA-sequencing tools presents applied scientists with a double-edged sword, as highlighted by a recent Nature paper (Dance 2022): a wealth of choices for data analysis, yet an overwhelming challenge in navigating through an ever-growing and complex array of methodologies.

To address the complexities of selection among the multitude of single-cell RNA-sequencing tools, the research community has placed a considerable emphasis on benchmarking. This is evident not only through numerous individual efforts to publish benchmark papers on various topics (You et al. 2021; Luecken et al. 2021; Mereu et al. 2020; Tian et al. 2019; H. Li et al. 2023) but also through the emergence of community-focused initiatives, such as the “Open Problems in Single-Cell Analysis” (https://openproblems.bio/), a web portal for hosting various single-cell analysis tasks such as cell-cell communication and spatial decomposition.

In the broader bioinformatics community, qualitative guidelines have been suggested to ensure high quality benchmarking. Notably, Weber and colleagues (Weber et al. 2019) proposed ten essential guidelines based on their experiences in computational biology. Further, Mangul’s group (Mangul et al. 2019) reviewed 25 studies across various topics in computational omics research and proposed principles that enhance the reproducibility and transparency of results. Currently, it remains unclear to what extent various research communities have embraced established benchmarking practices. In light of the rapid advancement of methodologies and the acknowledged significance of benchmarking within the single-cell research field, this field stands out as a good exemplar for exploring the present state of benchmarking practices and helping to understand gaps that necessitate community attention. Recently, Sonrel and colleagues provided the first paper (Sonrel et al. 2023) that quantitatively reviewed a collection of 62 single-cell benchmarking papers. They emphasised the technical aspects of the single-cell benchmark works and highlighted the need for code reproducibility, interoperability and extensibility. However, due to the multifaceted nature of the single-cell field encompassing not only technical aspects but also other considerations such as methodology, dataset and biological context, there is a need for a more comprehensive evaluation that takes account of the wider aspects of the single-cell field.

To achieve a quantitative understanding of the current landscape in single-cell benchmarking, we embarked on an extensive review. This involved conducting a systematic literature search for single-cell papers published between 1/1/2017 to 29/8/2024 via a PRISMA strategy (Page et al. 2021). The final collection includes 282 papers. This includes every benchmark-only paper in single-cell research (n = 130), where each paper is read by two different readers at least, along with a set of 152 method development papers that incorporate benchmarking elements, representing a collective human effort over approximately 300 hours. We designed a two-stage survey with nine sections such as study design and downstream evaluation metric, each aligning with a crucial aspect of benchmarking to gather key information from each paper and asked participants in the research group to fill in the survey (Methods). In total, more than 70 variables were collected. Through this systematic approach, we provide a comprehensive overview of the current state of play in single-cell benchmarking and raise awareness among the computational community regarding the challenges and opportunities.

## Results

### A comprehensive study design to evaluate single-cell benchmarking studies

To examine the current landscape of single-cell benchmarking studies systematically, we employed a two-stage process. In the first stage, we designed a pilot study, where we carefully read 17 selected benchmarking studies and identified all evaluation strategies presented in those studies (Figure 1a). Next, we categorised all these strategies into nine separate sections representing information relating to datasets, methods, accuracy criteria, scalability, stability, downstream analysis, context specific discovery, communication and software (Figure 1b). We then designed a survey with multiple choices and open-ended questions that facilitate the capture of all nine aspects of evaluations associated with a given benchmarking study. The finalised survey was given to volunteer participants who then reviewed and recorded the information on each paper as survey response (Figure 1c).

**Figure 1.**
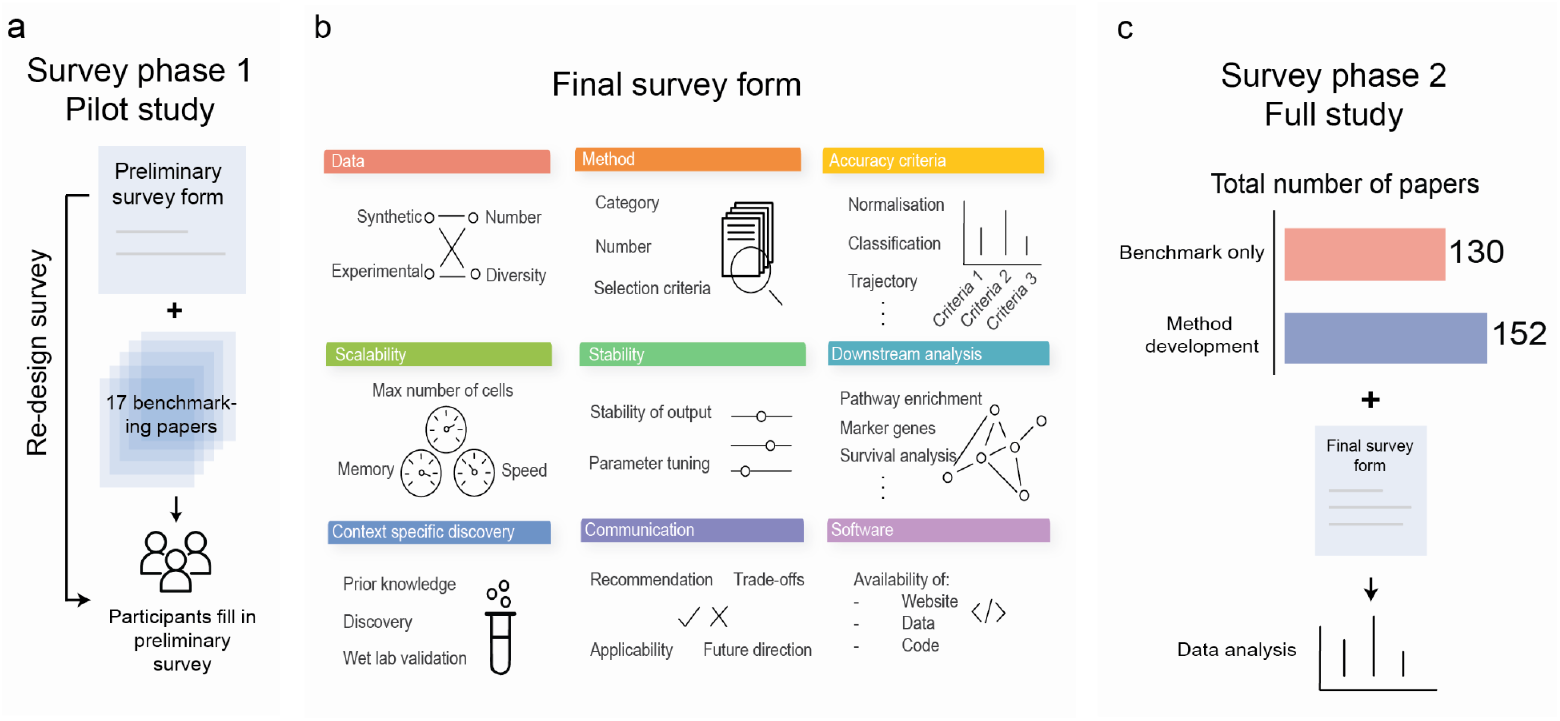
Schematic overview of the design of the survey. The survey was designed in a two-stage process. a. During the pilot study stage, a preliminary survey was designed that incorporated insights from 17 benchmarking papers. The survey was distributed to participants to collect feedback for the final survey in an iterative form. b. The final survey covered nine categories of evaluation, including data, method, accuracy criteria, scalability, stability, downstream analysis, context specific discovery, communication and software. c. In the full study stage, the participants reviewed a total of 282 papers, 130 from benchmark-only papers and 152 from method development papers, which provided the data for the analysis of this study.

We performed a systematic literature search using key terms such as “single-cell”, “systematic evaluation”, “benchmark” (Supplementary Figure 1, 2). We followed the PRISMA flow diagram to systematically record the selection process, including the inclusion and exclusion criteria. We intentionally included both benchmark-only papers (BOP) and method development papers (MDP) with a benchmarking component. We acknowledge that method development papers often include a benchmarking section to illustrate the effectiveness of their proposed methods. By including both the BOP and MDP papers, we are able to present a more comprehensive overview of the benchmarking practice in single-cell research.

In total, 33 readers contributed to the reading with a total of 433 survey responses (Supplementary Table 1). This corresponds to a total of 282 unique papers including 130 benchmark-only papers and 152 method development papers (Supplementary Figure 3), where all benchmark-only papers were read at least by two readers and their consensus was taken. The collection of papers involves 13 different technologies including single-cell RNA-seq (scRNA-seq), single-cell genomics, single-cell ATAC-seq (scATAC-seq), single-cell multiomics, spatial transcriptomics (ST) and spatial imaging. The benchmarked topics are diverse ranging from initial analysis such as batch correction, and intermediate analysis such as cell annotation, to downstream analysis such as differential expression, as well as analysis pipelines and data (Supplementary Figure 4).

### The current landscape of benchmarking studies

We examined the landscape of benchmarking from benchmark-only papers and method development papers. Table 1 and Supplementary Figure 5 show an overview of the results from nine components of the survey. Figure 2 presents key criteria and the percentage of benchmark-only and method development papers that met each criterion. Interestingly, we observed a broadly similar landscape of characteristics between BOPs and MDPs (Figure 2a-b, R = 0.83), despite these papers serving distinct purposes for the research community. This similarity reveals that the general benchmarking challenges are common in our community from both a review and development perspective.

**Table 1:** Summary of key findings from the survey. For the criterion “downstream analysis”, we removed the papers classified under the broad category as “downstream analysis” in the calculation. The reason is that we are only interested if the paper performed downstream analysis if the method itself is not a downstream analysis method. For the criterion “recommendation”, we did not calculate this criterion for method development papers. The reason is that we assume method papers would recommend their own proposed method and this criterion is not relevant.

| Benchmarking aspects | Criteria | Explanation | Key findings for benchmark-only papers (BOP) | Key findings for method development papers (MDP) |
| --- | --- | --- | --- | --- |
| Paper category | Paper category | Number of studies in benchmark-only papers and in method development papers | N = 130 | N = 152 |
| Data | Types of data | The types of data the study used to perform method evaluation. Whether they used experimental dataset, or synthetic datasets, or both. | Experimental: 54 (42%)<br>Synthetic: 3 (2%)<br>Both: 73 (56%) | Experimental: 82 (54%)<br>Synthetic: 1 (1%)<br>Both: 69 (45%) |
| | Number of experimental datasets | Number of experimental dataset used (median ± sd) | 6 ± 30 | 5 ± 43 |
| | Number of synthetic datasets | Number of synthetic dataset used (median ± sd) | 12 ± 2306 | 6 ± 773 |
|  | Diversity of experimental data | Whether the experimental dataset is considered diverse (Papers that use synthetic datasets only are excluded from the counting) | Yes: 105 (83%)<br>No: 16 (13%)<br>Not sure: 6 (5%) | Yes: 125 (83%)<br>No: 22 (14%)<br>Not sure: 4 (3%) |
|  | Diversity of synthetic data | Whether the synthetic dataset is considered diverse (Papers that use experimental datasets only are excluded from the counting) | Yes: 49 (65%)<br>No: 20 (26%)<br>Not sure: 7 (9%) | Yes: 45 (64%)<br>No: 22 (31%)<br>Not sure: 3 (5%) |
| Methods | Number of methods | Number of methods benchmarked (median ± sd) | 10 ± 28 | 4 ± 4 |
|  | Inclusion of selection criteria | Whether the study mentioned about selection criteria of the methods | Yes: 63 (49%)<br>No: 62 (48%)<br>Not sure: 5 (4%) | Yes: 49 (32%)<br>No: 94 (62%)<br>Not sure: 9 (6%) |
| Accuracy | Variability of score | Whether the study shown variability of scores (for example, in terms of boxplot) | Yes: 95 (73%)<br>No: 34 (26%)<br>Not sure: 1 (1%) | Yes: 89 (59%)<br>No: 63 (41%) |
|  | Overall comparison | Whether the study shown overall comparison figure or table of all evaluated methods | Yes: 62 (48%)<br>No: 68 (52%) | Yes: 46 (30%)<br>No: 106 (70%) |
| Scalability | Speed measured | Whether speed of the methods was measured | Yes: 81 (62%)<br>No: 49 (38%) | Yes: 69 (45%)<br>No: 83 (55%) |
|  | Memory measured | Whether memory usage of the methods was measured | Yes: 38 (29%)<br>No: 92 (71%) | Yes: 32 (21%)<br>No: 120 (79%) |
| | Max number of cells | If either speed or memory usage was measured, what was the maximum number of cells tested (median ± sd) | 12,706 ± 349,684 | 33,611 ± 5,944,397 |
| Stability | Sensitivity analysis | Whether sensitivity analysis was performed, for example, by subsampling the data and examine the impact on performance | Yes: 47 (36%)<br>No: 83 (64%) | Yes: 37 (24%)<br>No: 115 (76%) |
|  | Tuning | Did the study performed parameter tuning of the methods, or used default parameter settings | Parameter tuning: 31 (24%)<br>Default setting: 99 (76%) | Parameter tuning: 22 (16%)<br>Default setting: 130 (86%) |
| Downstream | Downstream analysis | Whether the study examined downstream analysis using the output produced from the methods | Yes: 56 (58%)<br>No: 38 (40%)<br>Not sure: 2 (2%) | Yes: 64 (57%)<br>No: 47 (42%)<br>Not sure: 1 (1%) |
| Context specific confirmation/discovery | Prior knowledge | Whether the methods capture any prior knowledge | Yes: 24 (19%)<br>No: 101 (78%)<br>Not sure: 5 (4%) | Yes: 62 (41%)<br>No: 80 (53%)<br>Not sure: 10 (7%) |
|  | Discovery | Whether the methods claimed any new discovery | No: 130 (100%) | Yes: 33 (22%)<br>No: 111 (73%)<br>Not sure: 8 (5%) |
|  | Wet lab validation | Whether the study performed any wet lab validation | Yes: 4 (3%)<br>No: 126 (97%) | Yes: 2 (1% )<br>No: 149 (98%)<br>Not sure: 1 (1%) |
| Communication | Recommendation | Whether the study provided any recommendation of the best methods | Yes: 110 (85%)<br>No: 16 (12%)<br>Not sure: 4 (3%) | Not applicable |
|  | Applicability | Whether the study examined applicability of the methods, for example, certain methods being applicable for certain tasks | Yes: 70 (54%)<br>No: 55 (42%)<br>Not sure: 5 (4%) | Yes: 42 (28%)<br>No: 103 (68%)<br>Not sure: 7 (5%) |
|  | Trade-offs | Whether the study examined trade-offs of the methods | Yes: 65 (50%)<br>No: 59 (45%)<br>Not sure: 6 (5%) | Yes: 27 (18%)<br>No: 114 (72%)<br>Not sure: 11 (7%) |
|  | Future directions | Whether the study suggested any future studies | Yes: 75 (58%)<br>No: 53 (41%)<br>Not sure: 2 (2%) | Yes: 82 (54%)<br>No: 68 (45%)<br>Not sure: 2 (1%) |
| Software | Website | Whether the study produced a website hosting the results | Yes: 8 (6%)<br>No: 122 (94%) | Yes: 10 (7%)<br>No: 142 (93%) |
|  | Data availability | Whether apart from providing the accession code, the study provided the curated data available for direct download | Yes: 68 (52%)<br>No: 62 (48%) | Yes: 62 (41%)<br>No: 90 (59%) |
|  | Code availability | Whether the code or package is available | Yes: 97 (75%)<br>No: 33 (25%) | Yes: 136 (90%)<br>No: 16 (10%) |

**Figure 2.**
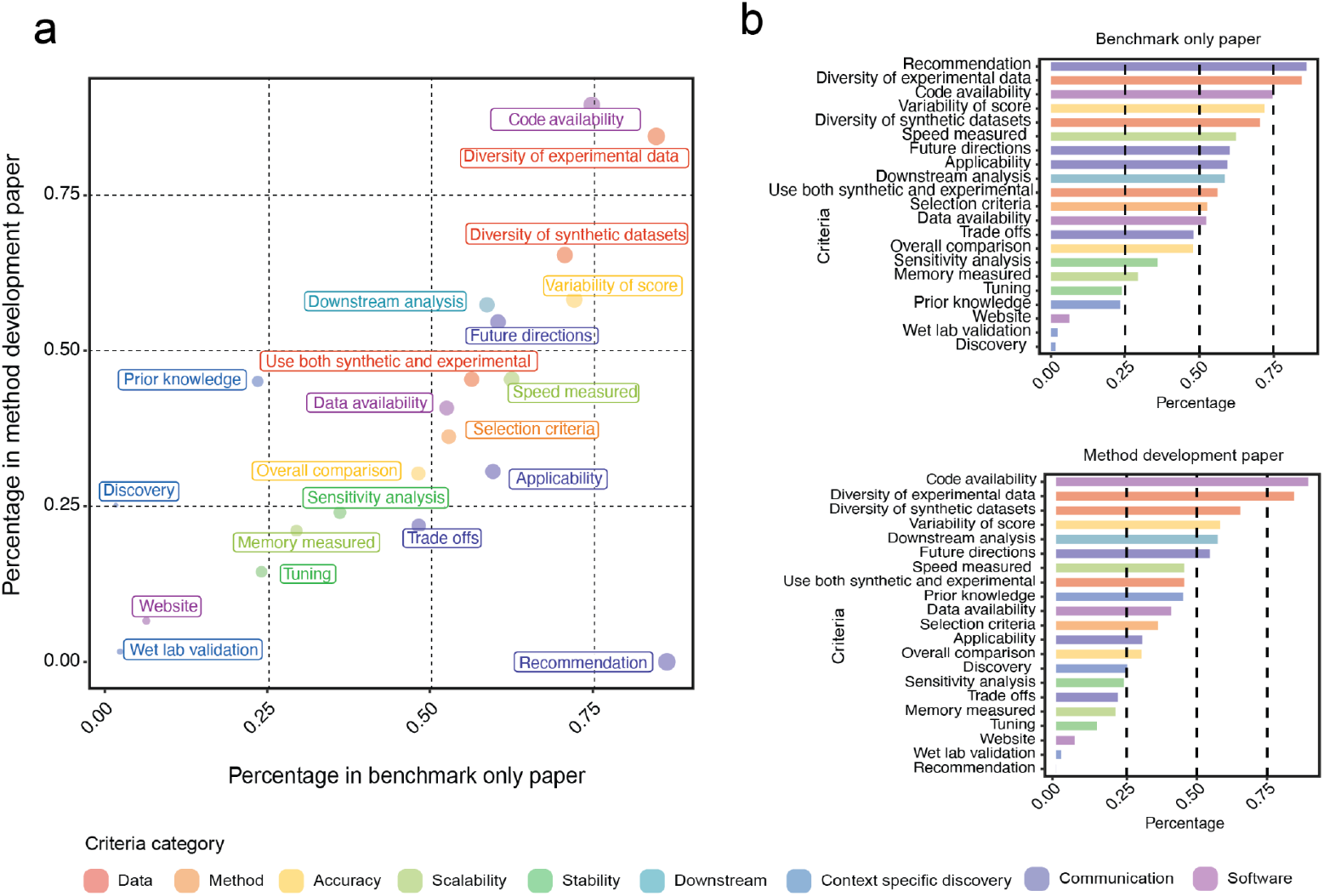
Percentage of criterion fulfilment. a. The percentage of papers that fulfilled each criterion. The x-axis indicates benchmark-only papers and y-axis indicates method development papers. Note that the “recommendation” criterion is considered not applicable for method development papers and is therefore given a score of 0. The criteria “downstream analysis” is calculated based only on the papers that belong to the analysis category of “data”, “initial analysis” and “intermediate analysis”. b. Ranking of the criteria in benchmark-only paper and new method development paper, ordered by the percentage of papers that fulfilled each criterion.

Delving into the specific evaluation criteria, we start with the two fundamental components of benchmarking: the methods chosen for benchmarking and the characteristics of the datasets to which these methods are applied. We observed that, on average, BOPs featured a more extensive evaluation of methods (median = 10 methods) compared to MDPs (median = 4 methods). However, it is worth noting that it is often unclear why certain methods were chosen, as less than half of all papers reported selection criteria of the methods (49% in BOP; 32% in MDP). Expectedly, almost all papers used experimental datasets (97% in BOP; 99% in MDP), with the median number of datasets used being similar across the two paper types (median = 6 in BOP; median = 5 in MDP). Synthetic datasets, often important in generating specific scenarios, are used more prevalently in benchmark-only papers (58% in BOP; 46% in MDP), with the median number of datasets also being higher in the benchmark-only paper (median = 12 in BOP; median = 6 in MDP).

### The performance of a method is multi-faceted and extends beyond accuracy measurement

We observed that more than half of the papers from both categories recognise downstream biological application as an important method assessment (58% in BOP; 57% in MDP). New biological discoveries were claimed by 22% of the method development papers (0% in BOP). The performance of a method can also be context specific and it is important to communicate the advantages and limitations of different methods. Applicability, which refers to the suitability of a method for a specific task, was conducted in 54% of benchmark-only papers and 28% of method development papers. Trade-off analysis, which refers to the compromise between different aspects of a method, was performed in 50% of benchmark-only papers and 18% of method development papers. Stability of methods is critical for reproducible research. However, we noted only a minority of papers performed sensitivity analysis (36% in BOP; 24% in MDP) such as assessing the impact of data subsampling on method performance. In terms of scalability, while a significant number of papers reported speed (62% in BOP; 45% in MDP), only a minority of papers measured memory usage (29% in BOP; 21% in MDP).

In terms of software availability, we observed the sharing of code is a common practice in the single-cell field with 90% of method papers providing code and 75% in benchmark-only papers. The provision of curated datasets and not just data accession ID is less common, with less than 52% of BOP and 41% of MDP fulfilling this criterion. Whilst it is not journal requirement for studies to provide interactive websites for accessing the results, we observed that eight studies in benchmark-only studies and ten studies in method development papers provided websites.

### The challenges of consistency in multiple benchmarking studies on a single topic

We observed a significant number of topics that have attracted multiple benchmarking studies. Across the 33 topics covered in the benchmark-only papers, 19 of them (58%) have more than one study (Supplementary Figure 3). Next, we focused on popular topics with at least five studies within the data type of scRNA-seq, such as dimension reduction (n = 10), cell type/state identification (n = 10) and differential expression (n = 8). We unexpectedly found a limited number of overlaps in the methods evaluated across the multiple studies (Figure 3a, Supplementary Figure 6a). For example, in the eight differential expression benchmarking studies, the majority of the methods (60%; 22/37) were only benchmarked in one paper.

**Figure 3.**
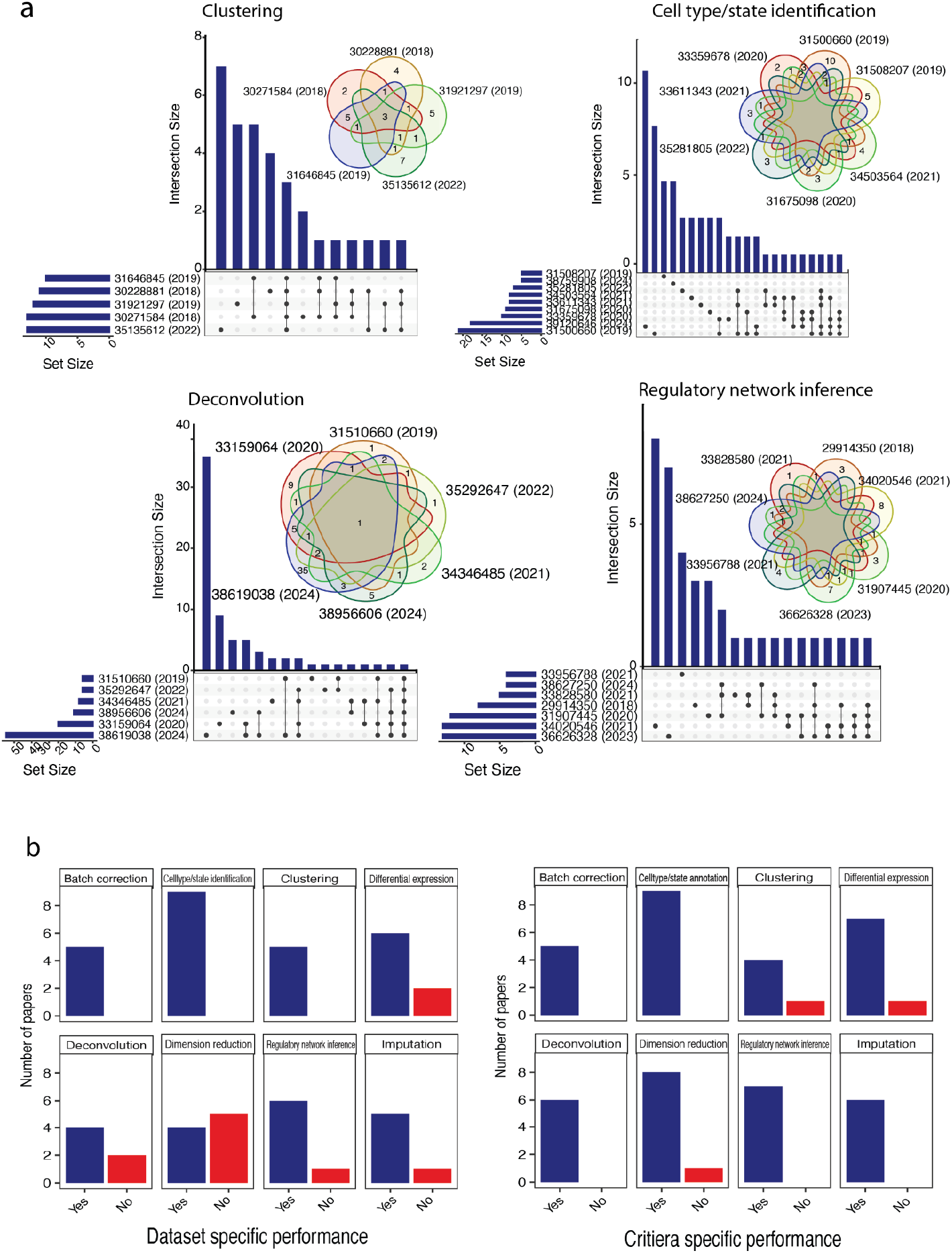
Inspecting consistency across multiple benchmarking studies. a. Both the upset plot and Venn diagram show the number of common methods evaluated across different benchmarking papers within each topic. Each paper is denoted by the PMID number. b. Number of papers reporting dataset-specific performance of methods. c. Number of papers reporting criteria-specific performance of methods.

The lack of overlap in the evaluated methods raises the question of how to best consolidate the knowledge across multiple benchmarking studies. Is it even possible to derive consensus rankings for the methods? In parallel, effective consolidation is further hampered by the different criteria and datasets used across the benchmark. Almost all benchmark papers examined (96%) reported criteria-specific performance, while 80% of benchmark papers reported dataset-specific performance (Figure 3b,c). The criteria and datasets used by each benchmarking paper are different with less than 10% overlap in many of the topics (Supplementary Figure 7, 8). For example, one might expect accuracy of cell type classification to be a key criterion for all cell type/state identification papers. We found each of these papers examined different measures of accuracy such as overall accuracy and microF1, as well as different scenarios such as the effect of using different sizes of reference data, the impact of feature selection and the impact of batch correction. The combination of different scenarios and different measures resulted in only three out of the total of 83 criteria (4%) being used in more than one paper among the ten cell type/state identification papers. While the emphasis on criteria and dataset-specific evaluations is valuable, a key question arises: how can we effectively synthesise knowledge from multiple benchmarking studies to enhance our understanding?

### Data diversity for comprehensive method evaluation

Next we focused on the choice of datasets in benchmarking evaluations. Surprisingly, the median number of experimental datasets used in benchmark-only papers (6 ± 30) is comparable to that observed in method development papers (5 ± 43) (Table 1, Figure 4a), with P-value = 0.29 under Welch Two Sample t-test. This observation poses questions on whether the current benchmarking studies utilise enough datasets for a comprehensive assessment, as method performance can vary across different datasets.

**Figure 4.**
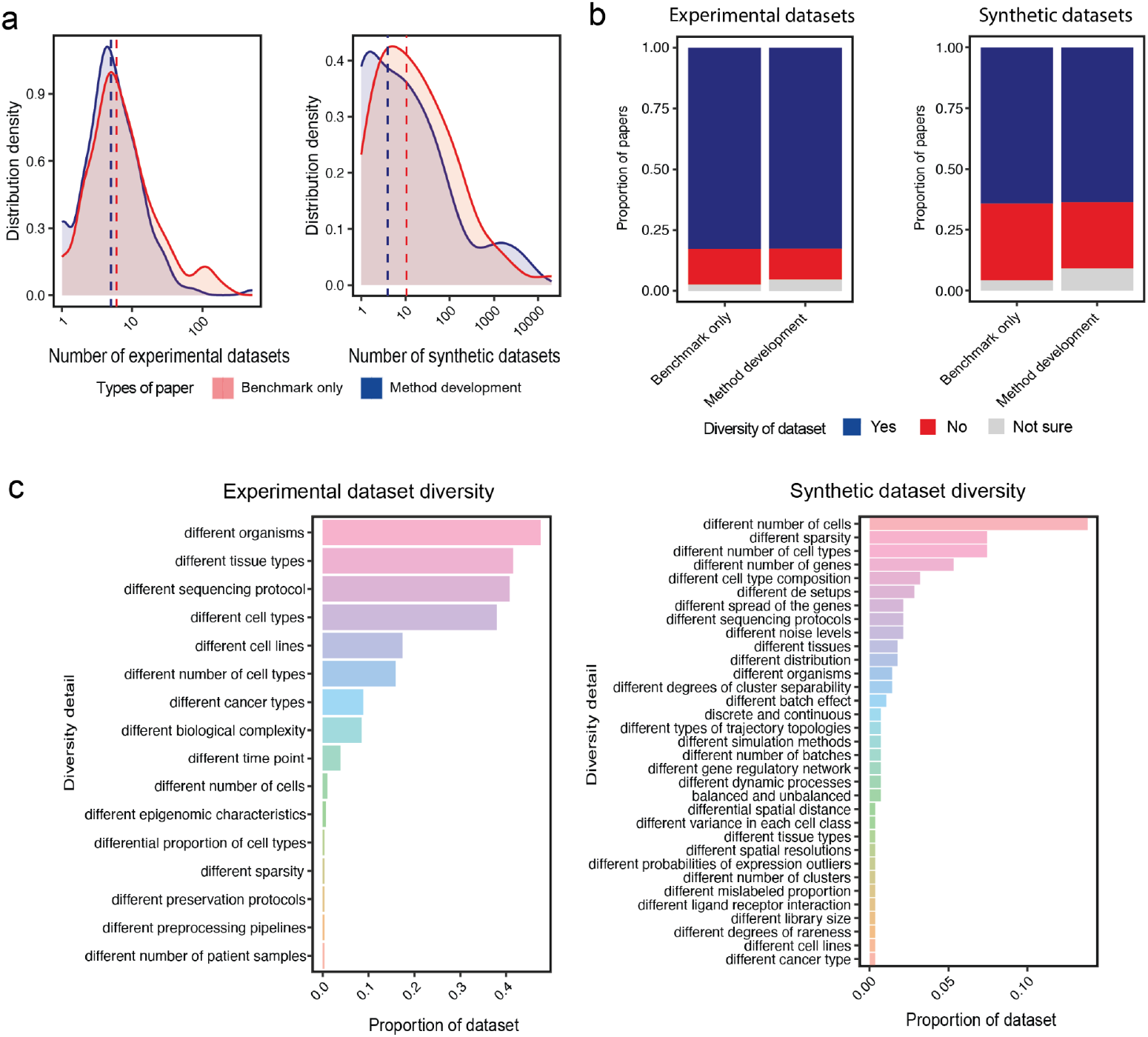
Usage of datasets in benchmark-only and method development papers. a. Distribution of the number of experimental and synthetic datasets in benchmark-only and method development papers. Vertical line indicates the median number of dataset used. b. Proportion of papers with experimental and synthetic datasets that are considered to be diverse. c. Detail of the diversity in experimental and synthetic datasets, ordered by the proportion of the datasets in each diversity category. The total proportion may exceed 1 as one dataset can exhibit multiple types of diversity.

In addition to the number of datasets, the diversity of datasets, both experimental and synthetic, is also an important factor for method evaluation. We found, irrespective of the type of papers, a greater percentage of readers indicated that the experimental data used were diverse (83% in both BOP and MDP), compared to synthetic data (65% in BOP and 64% in MDP) (Table 1) (Figure 4b). One possibility is that despite a study generating multiple synthetic datasets, they could have been derived from a limited set of experimental data. However, synthetic data exhibited a broad range of diversity. In experimental datasets, the diversity predominantly arose from various characteristics inherent to the dataset, such as sequencing protocol and cell/tissue type (Figure 4c). In contrast, the diversity in synthetic datasets can be carefully designed to encompass specific scenarios that are not easily accessible in experimental datasets such as a spectrum of sparsity levels. Incorporating both experimental and synthetic datasets can offer a more comprehensive understanding of method performance across diverse biological scenarios.

### Unravelling temporal trends in benchmarking practices

We sought to understand how the landscape of benchmarking studies changes over time (2017-2024) among our 282 papers by using numeric scores to represent selected criteria (Supplementary Table 2) where a higher score indicated the paper has fulfilled a greater number of criteria. We recognise that the year when a field first emerged would confound criteria such as the number of methods evaluated in each year, and thus, we adjusted publication year into a relative publication year based on when the field first emerged (see Methods) (Figure 5a).

A notable finding is the trend of the number of methods evaluated across adjusted publication years and the emerging challenge of ‘benchmarking fatigue’. The overall scores of papers as well as many of the individual criteria such as the number of datasets used remained stable over time (Figure 5b, Supplementary Figure 9). In contrast, both the benchmark-only papers and method development papers revealed an increasing trend in the number of methods compared across the years (Figure 5c). As the relative publication year progressed from year 0 to year 9 within a field, the median number of methods evaluated increased from 4 to 13 in benchmark-only papers. Whilst the increasing number of methods reflects active development of the single-cell field, it also motivates consideration of how to approach the ever-increasing number of methods in terms of evaluation. Furthermore, we noted that 40% of BOP and 50% of MDP evaluated more methods in their published versions compared to their bioRxiv counterparts, irrespective of the number of methods initially benchmarked in the bioRxiv version (Figure 5d), further adding to this benchmark fatigue.

**Figure 5.**
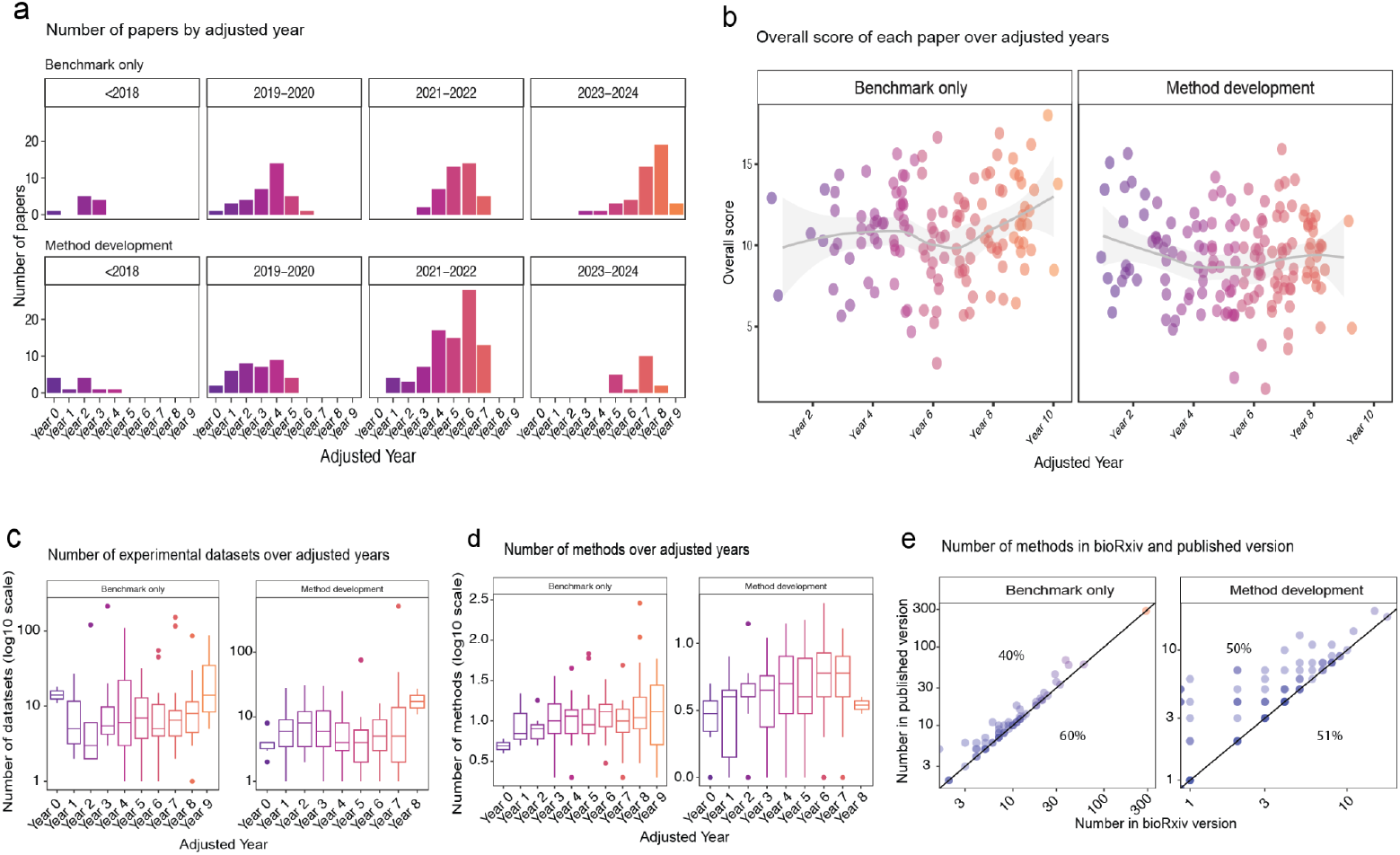
Temporal trends of selected benchmarking practices. a. Number of papers in adjusted publication year, stratified by their actual publication year. b. Overall score of each paper over adjusted publication years. c. Number of experimental datasets used in each paper over adjusted publication year. d. Trend of number of methods evaluated in each paper over adjusted publication year. e. Number of methods evaluated in bioRxiv and published versions of the papers. The percentage above the diagonal line indicates the proportion of papers where the number of methods is greater in the published version. The percentage below indicates the proportion of papers where the number of methods is equal in the published and bioRxiv version.

## Discussion

This study performed a systematic literature search and analysed a total of 282 papers in the single-cell literature space, consisting of 130 benchmark-only papers and 152 method development papers. It extensively examines nine different aspects of evaluation, including but not limited to the type of data used in the studies, number of methods benchmarked, accuracy criteria, stability aspects, downstream analysis, capture of context specific knowledge, communication aspect and the availability of resources. With the evaluation of these papers, we uncover various challenges and opportunities for the community to consider.

We identified multiple areas for further improvement. For example, less than half of the assessed papers reported selection criteria explaining why certain methods were included in the benchmark. This underscores the need for greater transparency in reporting. We noted only over 30% of BOP and over 20% of MDP papers in both paper categories assessed stability of method performance. Stability is one of the three major principles underlying data science, together with predictability and computability (Yu and Kumbier 2020). Unstable methods create confusion and irreproducible results in research. In terms of scalability, the evaluation of memory usage is an area requiring attention. As the size of single-cell data continues to surge, with datasets reaching millions of cells and atlases containing tens of millions of cells (M. Li et al. 2021; Rozenblatt-Rosen et al. 2017)), memory is becoming a key consideration in method selection. Interestingly, we note that while the availability of data is often required in many journals, almost less than half of studies provide processed data, with the rest of studies providing either no data or links to raw data. Data curation is a time-consuming step in research and the accessibility and reusability of the data as outlined by the FAIR Data Principles will benefit the scientific community (Wilkinson et al. 2016).

We noted several challenges in the current single-cell benchmarking field that necessitate a joint community effort. One of the questions is how to deal with the information presented in multiple benchmarking studies when each study examined a different collection of methods and datasets. Relatedly, the large proportion of studies that report dataset-specific method performance presents another challenge for applied researchers wanting to choose a method for their own datasets. To address these challenges, approaches for performing meta-analysis or a system for combining the results from multiple studies are needed. Once large-scale results are curated, they can be used to construct a dataset-specific recommendation system (Fang, Selega, and Campbell 2024) to pinpoint the potential methods for a given dataset.

However, the above approaches are not without their challenges. In the current single-cell benchmarking field, the input and output of the methods are rarely made available and researchers who want to build from existing benchmarks would need to reconstruct the benchmark from scratch (Sonrel et al. 2023). This situation calls for a collective effort from all researchers for more transparency in result sharing and for the development of novel approaches to create a consensus benchmarking framework that can be extended from different methods, datasets and criteria. To take this one step further, the single-cell community could establish a benchmarking consortium to define a set of guiding principles for future benchmarking studies, including the deposition of data and results. In the clinical field, for instance, the famous Cochrane Collaboration was established to provide continuing guidelines and advice on systematic review (Higgins et al. 2011) and is considered as a gold standard in the field.

We found that the median number of methods benchmarked in a study increases as the field progresses. While this reflects the natural progression of the field, it can lead to challenges such as benchmarking fatigue. A potential solution is a web-based system that holds existing benchmarking results and allows researchers to incorporate new methods into the existing results. This concept of “living benchmark” ideas has been adopted in domains such as microarray probeset summary (Cope et al. 2004) and scRNA-seq simulation methods (Cao, Yang, and Yang 2021). Notably, the “Open Problems in Single Cell Analysis” website (https://openproblems.bio/) hosts a similar platform for several popular areas such as batch correction and cell type annotation. Nevertheless, the number of datasets currently evaluated by the platform is limited, with typically around three datasets in each area. Besides including more datasets, the system should ideally enable new metrics or questions to be incorporated and existing methods to be c evaluated on the new metric. By joining community efforts, we believe such a system linking multiple datasets with metrics and questions can effectively allow new benchmarking studies to build on top of existing studies. This not only alleviates benchmarking fatigue by avoiding repeated evaluation of methods that have been previously assessed, but also facilitates integration of multiple benchmarking studies.

Finally, as large language models (LLMs) such as ChatGPT demonstrate ground breaking capability in textual interpretation, this collection of 433 survey responses curated by about 300 hours of human effort is a unique resource to adapt LLMs or assess LLMs on such task of identifying and processing key information from scientific literature. Many of the variables such as number of methods and number of datasets can be used as a basis for quantitative assessments.

In conclusion, this extensive assessment of a collection of 282 papers not only enhances our understanding of the current state of play in single-cell benchmarking but also underscores the need for collective efforts within the bioinformatics community. The survey result, made publicly available, is a valuable resource for the community for further investigation. We hope this work sets the foundation and raises a call for collective action from the single-cell community to set guidelines and advance benchmarking practices. We envisage the formation of a benchmarking consortia or collaborations similar to the Cochrane Collaboration will bring together experts from various single-cell areas and enhance the future of single-cell benchmarking.

## Methods

### Design of the study

#### Survey Design

To have a structured way of collecting information on each article, we designed a survey to contain multiple-choice and open-ended questions related to various aspects of journal articles. Each question was accompanied by a detailed explanation to ensure the clarity of the question and the quality of the response. This survey construction was achieved in a two-stage approach.

#### Stage I - pilot study

Stage I is a pilot study. Prior to the full-scale data collection, a pilot study was conducted with all authors. Authors read 17 single-cell papers published in renowned journals such as Nature Biotechnology, including both benchmark-only papers and method development papers to gather the metrics used in these papers. This stage is an iterative process where multiple meetings were held to discuss the questions included in the survey and update the survey. The final survey form included nine sections on various aspects including data, method, accuracy criteria, scalability, stability, downstream analysis, context specific discovery, communication and software. For a selected series of questions, that is, diversity of experimental dataset, diversity of synthetic datasets, types of downstream analysis performed, the questions were initially designed for users to input open text. Manual text analysis was then used to narrow these into categories and convert the questions into multiple choices, such that it is both easier for the reader to respond and easier for data cleaning. Users also had the option to type their responses for responses not included in the multiple choices.

#### Stage II - full study

In Stage II we performed the full study. We gave a 1-hour presentation session to the research group before people contributed, to ensure that people fully understood the questions. To ensure the survey result is not solely driven by the opinion of the authors of this study, we invited members, who are mostly PhD students, in our group to voluntarily contribute to the survey. For all benchmark-only papers, we ensured at least two readers for each paper to counter any potential error or bias. Details for addressing any inconsistency between two or more readers were given in the data processing Methods sections.

### Systematic selection of papers

#### Literature search

To perform a systematic literature search of papers, we used the following key index terms including “single cell”, “single-cell”, “spatial transcriptomics”, “benchmark”, “benchmarking”, “systematic evaluation”, “systematic comparison”, “comprehensive evaluation” and “review” to search on PubMed. We restrict the time of publication to between 1/1/2017 to 29/8/2024. This results in a total of 845 papers that were published since 2017. We manually screened all records to identify the relevant single-cell related papers and excluded any pre-print papers.

#### Collection of benchmark-only papers

In addition to the 845 papers, we identified 62 papers from the research conducted by Sonrel et al. (Sonrel et al. 2023) and a further three benchmark-only papers were recommended by readers. After excluding duplicates, this produced 885 papers. We then applied our filtering criteria on the time of publication and removed any pre-print papers on this collection. Subsequently, each paper underwent a screening process, leading to a final collection of 130 single-cell benchmark-only papers.

#### Collection of method development papers

From the 845 papers, most of the remaining papers that do not belong to benchmark-only papers are papers that proposed a new method. We randomly sampled 102 papers to be read as method development papers. We also received 51 recommendations from the readers. This resulted in a final number of 153 method development papers included in this study. We refer to this collection of papers as method development papers.

### Data processing and statistical analysis

#### Data cleaning and processing

We performed data cleaning on the survey response in order to address errors and inconsistencies. In particular, for all benchmark-only papers, each paper was read by at least two readers and inconsistency could arise as a result. We examined the columns corresponding to factual information and identified inconsistent responses. These columns are “Paper category”, “Memory measured”, “Speed measured”, “Website”, “Data availability”, “Types of data”, “Methods compared”, “Code availability”, “Number of experimental datasets”, “Sensitivity analysis”, “Tuning”, “Number of synthetic datasets”, “Max number of cells”, and “Overall comparison”. Another reader then manually cleaned the inconsistent values by reading the paper as well as the response by the two readers. Invalid responses were removed.

We also ensured responses in each column were consistent with one another. For example, if the answer to “Types of data” was “Both experimental and synthetic datasets” were used for evaluation, we ensured the “Number of experimental datasets” and “Number of synthetic datasets” were filled with the corresponding information from the paper. The final cleaned survey response is publicly available at https://github.com/SydneyBioX/sc_bench_benchmark as a CSV file.

#### Curation of additional variables

To extend the type of analysis possible, we manually curated further variables of interest. The following variables are created by reading the abstract of each paper: 1) the data type, ie, the types of single-cell omics data, 2) the broader topic type, we classified the papers into the following five categories: data, initial analysis, intermediate analysis, downstream analysis and analytical pipelines, 3) the finer topic type, we defined the specific task or purpose of the papers, such as classification, clustering, doublet detection, etc. Additionally, where there is a bioRxiv version of the published papers, we curated the number of methods compared in the bioRxiv version.

#### Temporal analysis

To quantitatively assess the trends in the benchmarking landscape across years, we transformed specific criteria into the range from 0 to 1 (Supplementary Table 2). The scores from each criterion were aggregated to generate an overall score for each paper, where a higher score indicates a greater fulfilment of the criteria.

We noted that certain criteria such as the number of methods compared can be confounded by the time when the field thrives. For example, it is not fair to compare the number of methods benchmarked in a spatial deconvolution paper with the number of methods benchmarked in scRNA-seq deconvolution papers despite them both being published in the same year. Spatial omics were introduced at a much later time than scRNA-seq and the number of methods available differ significantly.

To account for this confounding effect, we converted the publication year into what we termed as “dynamic publication year”. In detail, we utilised the single-cell RNA-seq tool website (Zappia and Theis 2021) which collects the publications on single-cell tools, the categorisation of the tools and the publication year. We calculated the year when each category had at least 5 papers, which we termed as year 0. We then subtract the publication year of each paper by their corresponding year 0 to create the dynamic publication year. For example, for a topic with five papers published in 2020, a benchmarking paper published on that topic in 2023 is then considered as being published in year 3. This thus effectively adjusts for the difference in time when each field was introduced.

## Backmatter

### Code and data availability

Code and anonymised survey result for reproducing the results of this study is publicly available at https://github.com/SydneyBioX/sc_bench_benchmark.

## Acknowledgement

This work is supported by the tremendous collective effort of members of the Sydney Precision Data Science Centre and WEHI. In total, 33 readers contributed to the data collection with a total of 433 readings. The authors would also like to thank other members at The University of Sydney, Sydney Precision Data Science Centre, School of Mathematics and Statistics, and Charles Perkins Center for their intellectual engagement.

