## Supplementary figures and tables for "The current landscape and emerging challenges of benchmarking single-cell methods"

### Supplementary Material

### Supplementary Figures

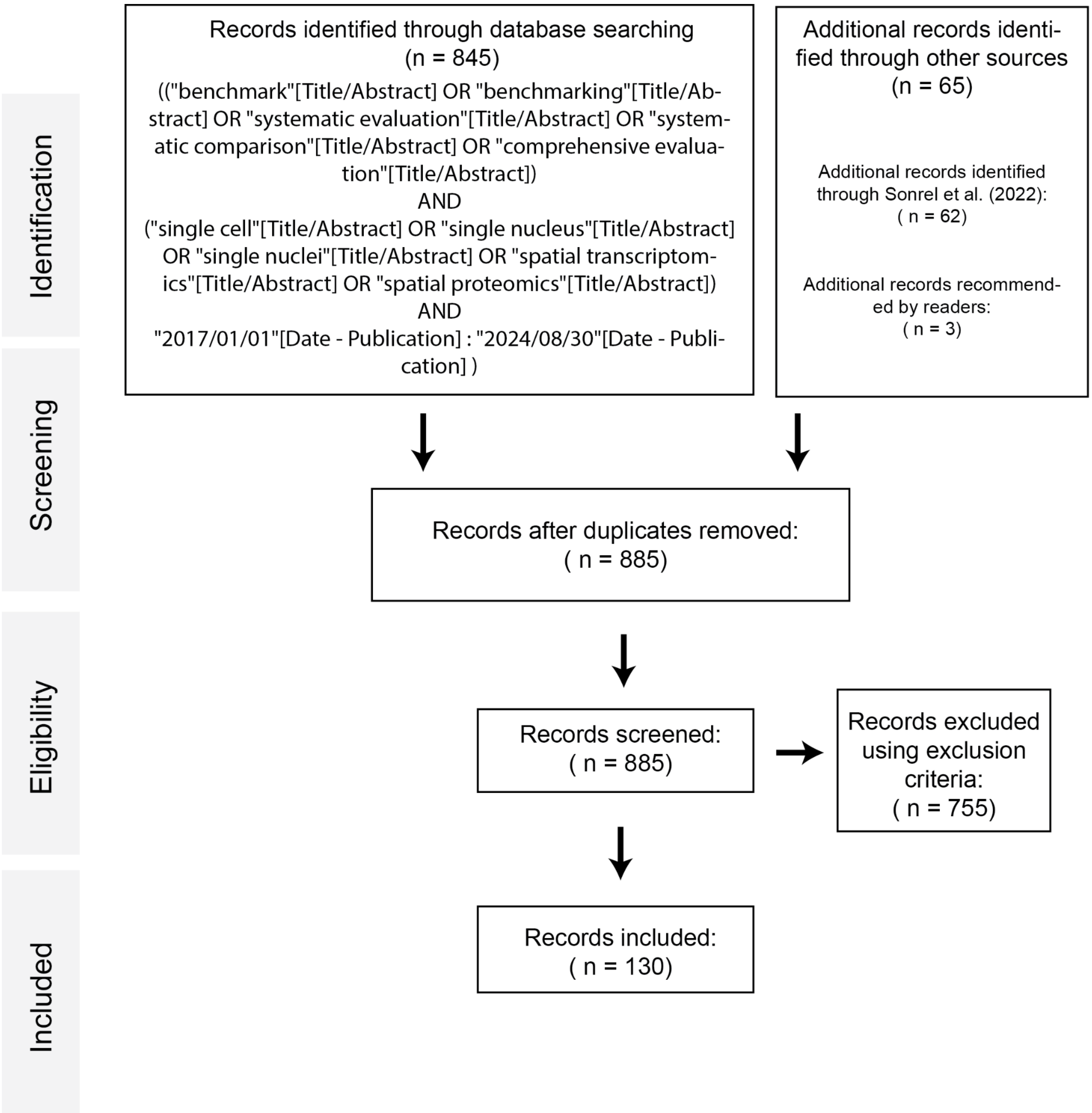


**Supplementary Figure 1. Flowchart for systematic literature search of benchmark-only papers.**
Using five combinations of key terms for PubMed searches and multiple rounds of inclusion and exclusion procedures, we identified a total of 95 benchmark-only papers.

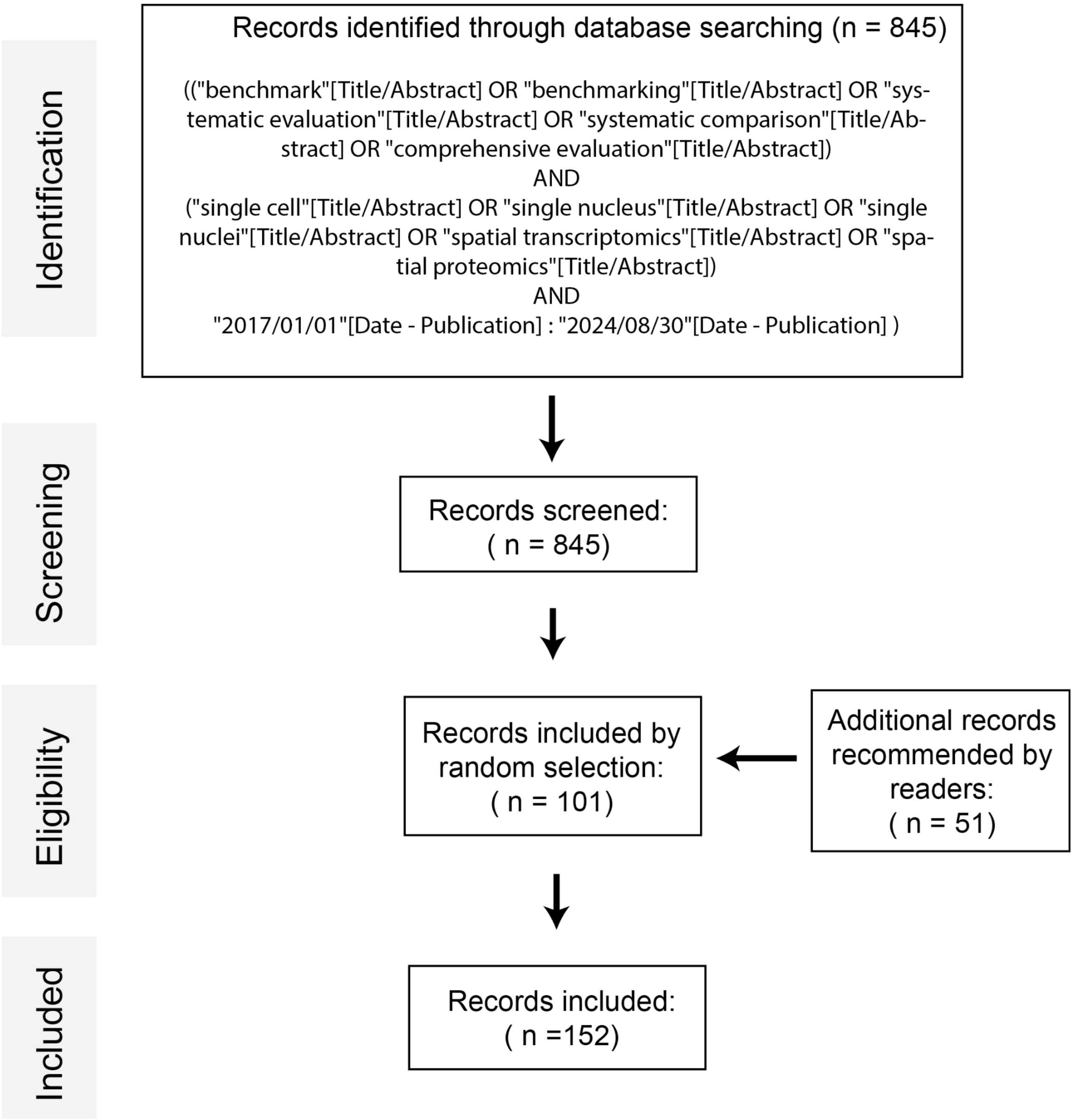


**Supplementary Figure 2. Flowchart for systematic literature search of method development papers.**

Using five combinations of key terms for PubMed searches and random selection and reader recommendation, we identified a total of 152 method development papers.

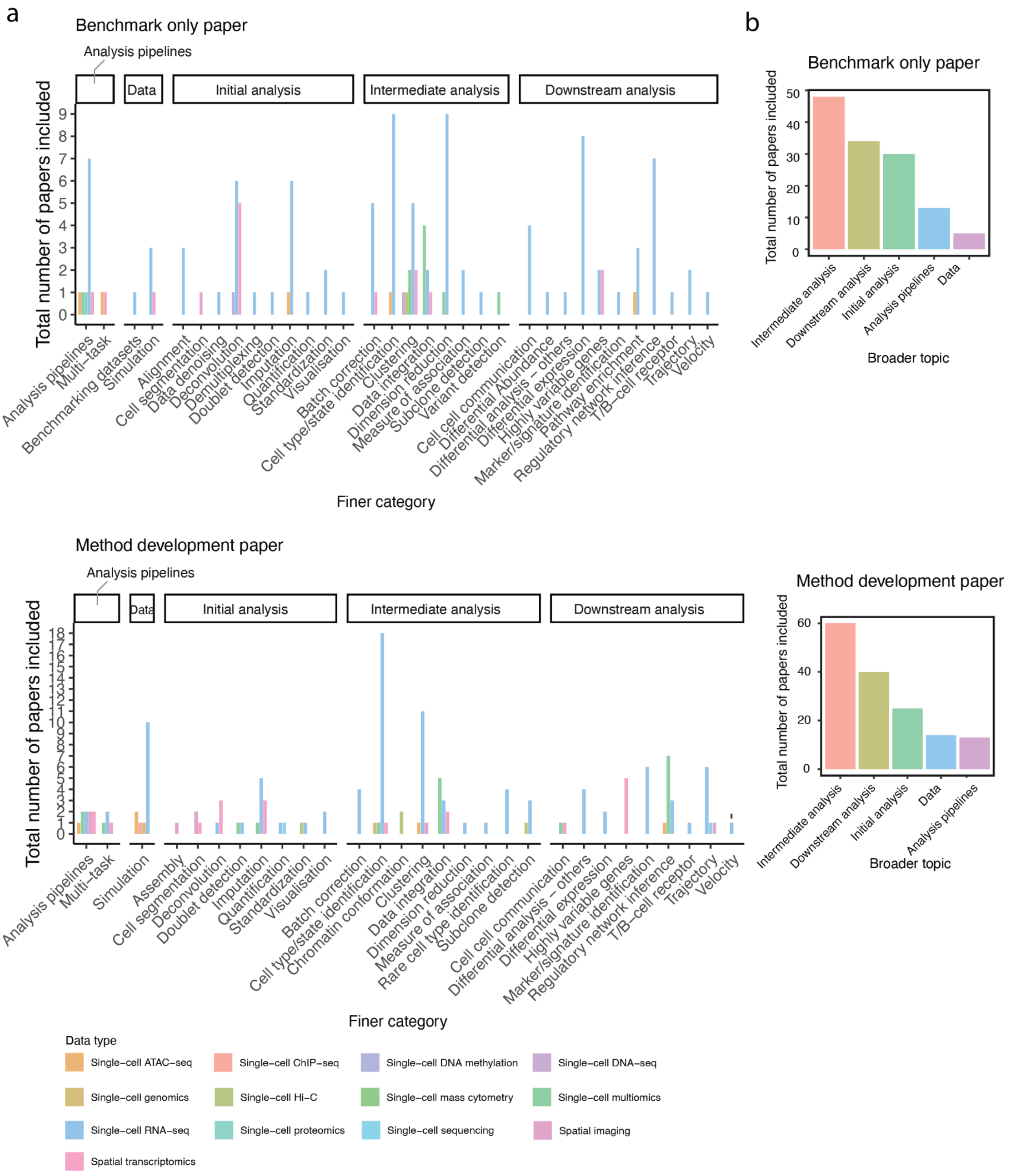


**Supplementary Figure 3. Distribution of the papers across finer and broader categories.**

a. Total number of benchmark-only and method development papers included in this study in each finer category, stratified by the broader category.

b. Total number of benchmark-only and method development papers in each broader category.

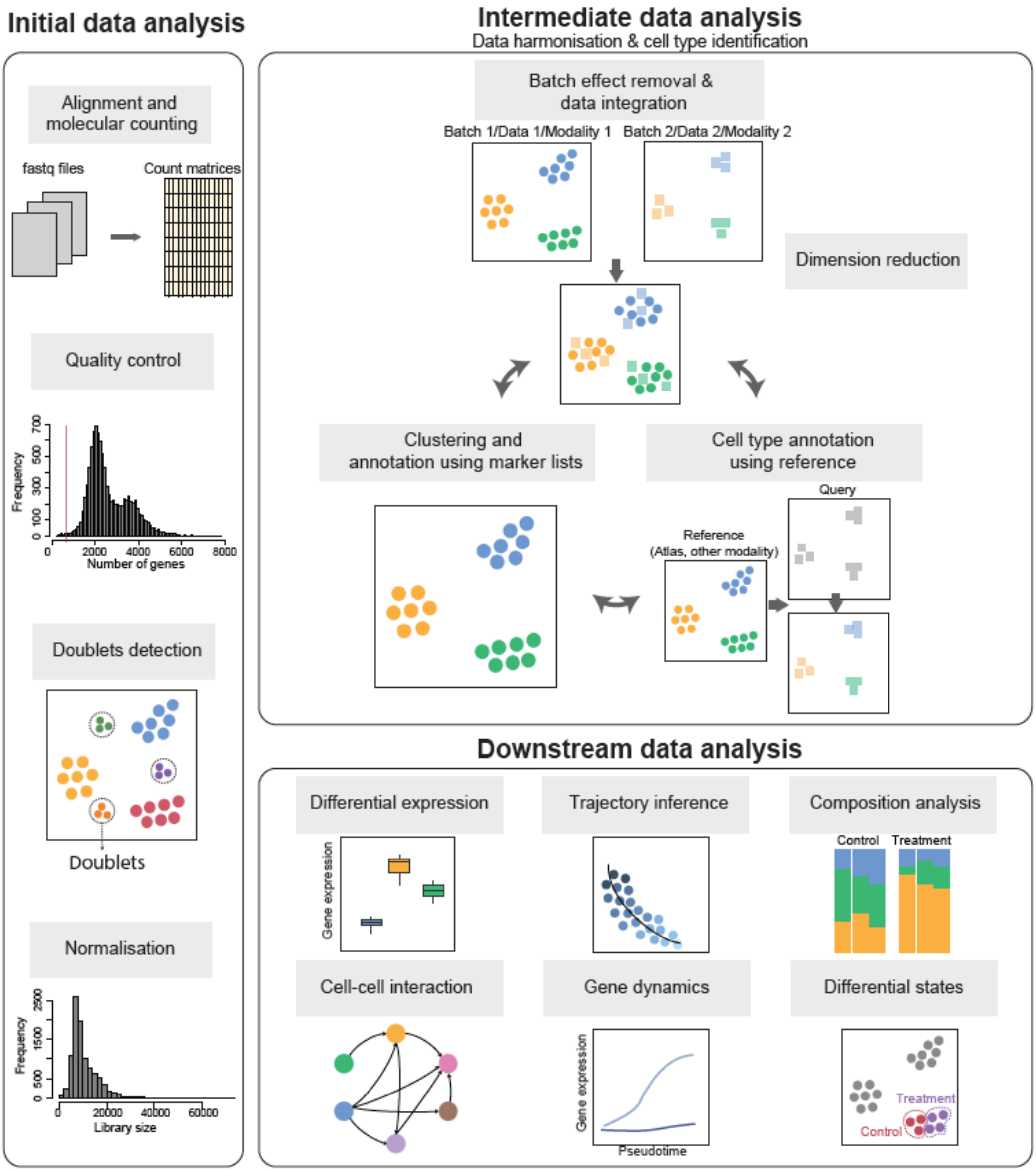


**Supplementary Figure 4. Examples of initial data analysis, intermediate analysis and downstream data analysis.**

In the study we classified the topic of each paper under five broad categories of “data”, “initial analysis”, “intermediate analysis” and “downstream analysis”. This figure gives selected examples for some of these categories.

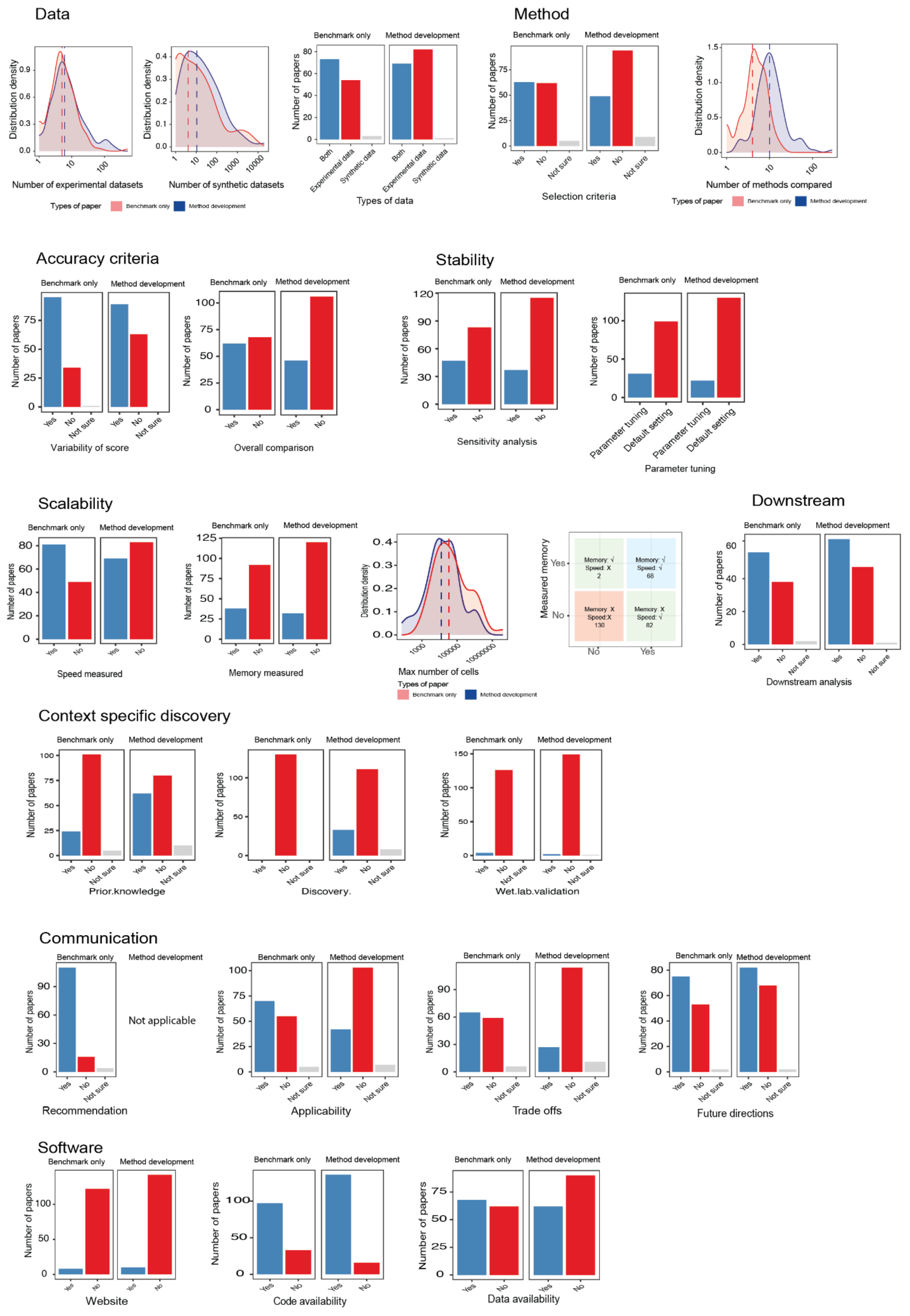


**Supplementary Figure 5. Overview of survey response.**

This figure provides a visual overview of the survey responses across the nine criteria categories.

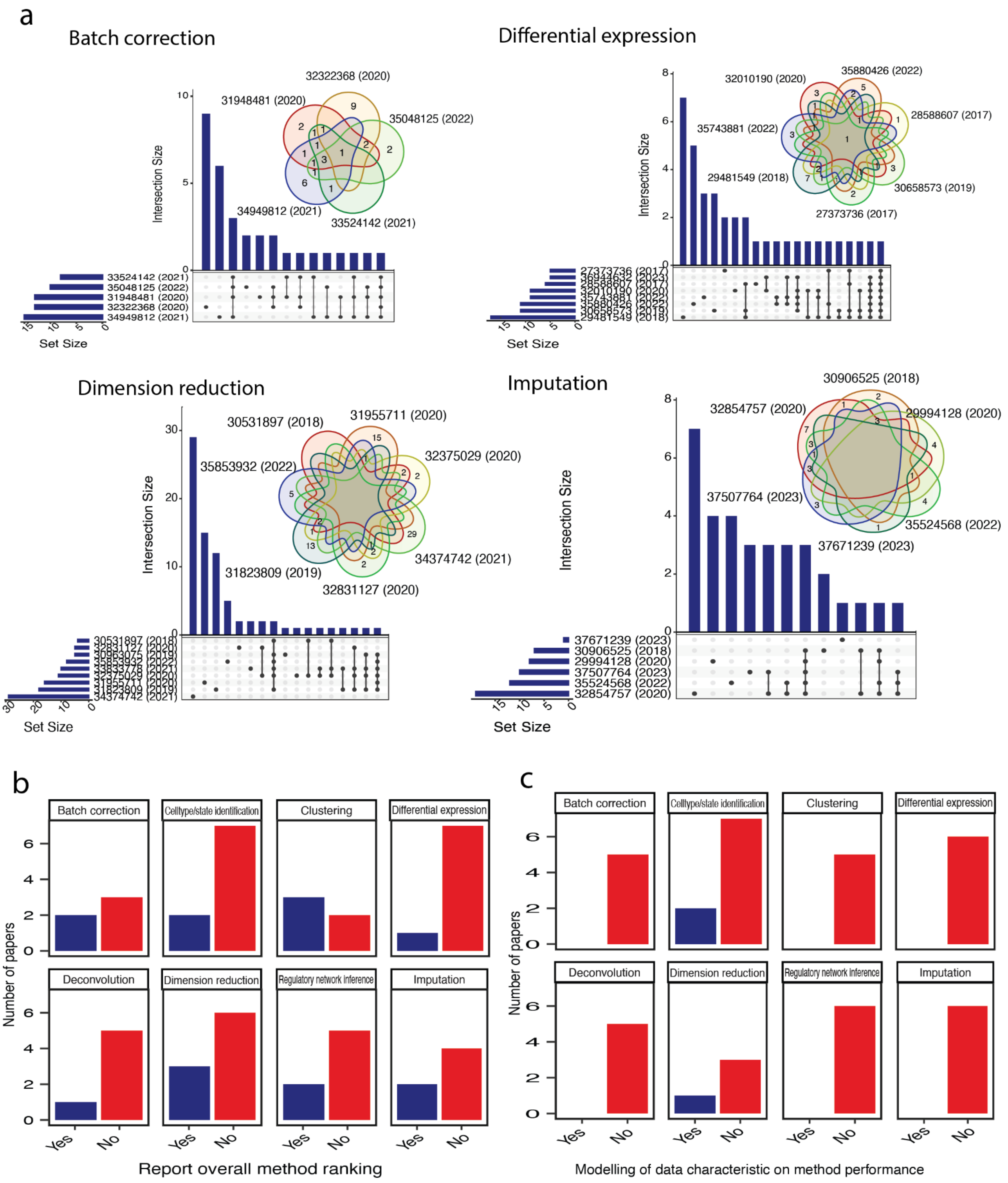


**Supplementary Figure 6. Inspecting consistency across multiple benchmarking studies.**

Topics with more than three benchmarking papers were assessed. a. Venn diagram and upset plot illustrate the common methods evaluated across multiple benchmarking papers within each category. Each paper is denoted by their PMID. b. Number of papers that report the overall method ranking in each category. c. Number of papers that analysed the impact of data characteristics on method performance in each category.

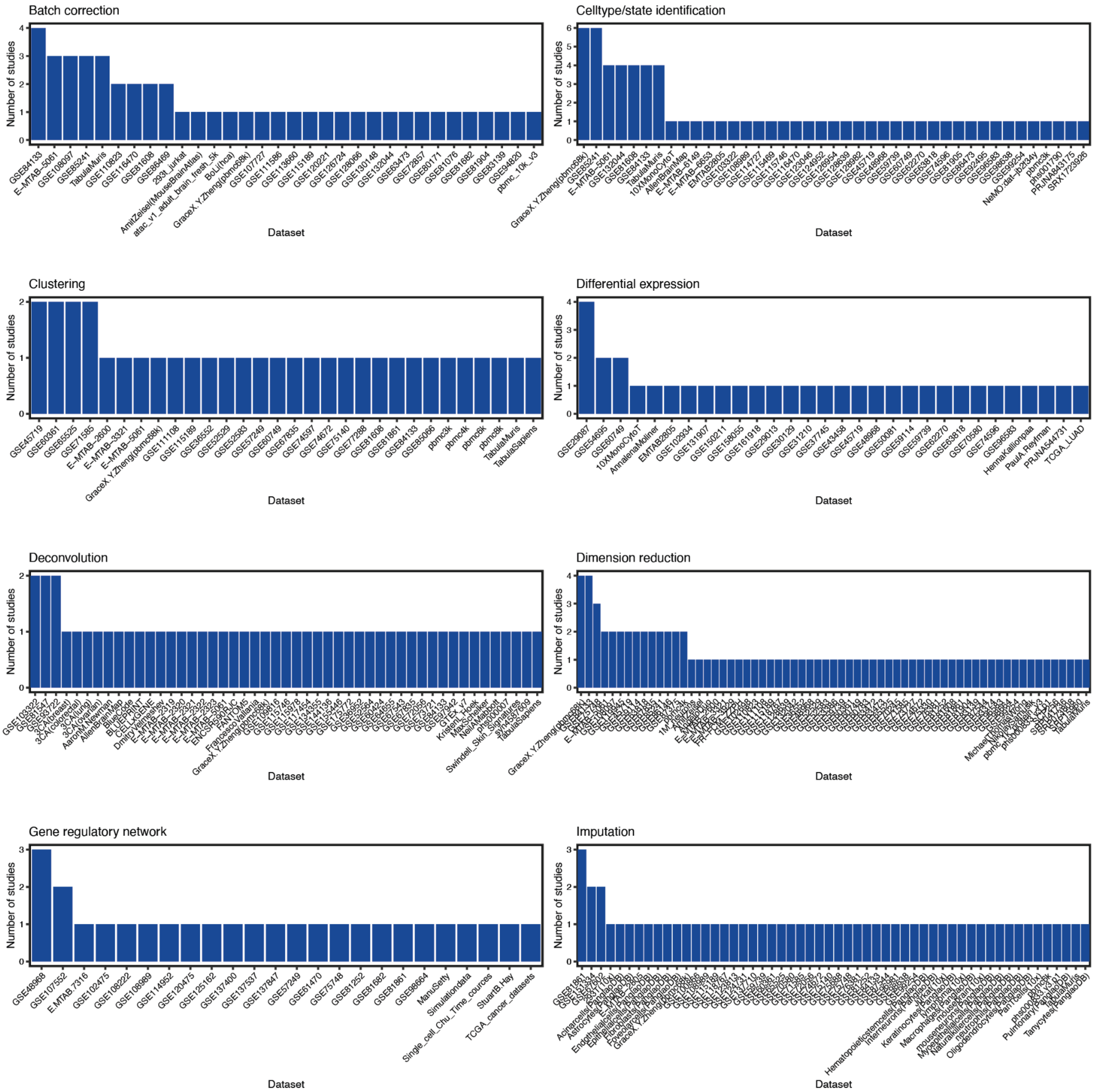


**Supplementary Figure 7. Overlap of the datasets used by multiple benchmarking studies.**

Topics with more than three benchmarking papers were assessed. The barplot denotes the number of papers that the datasets appeared in. Each dataset is denoted by their ID, such as the GEO accession ID.

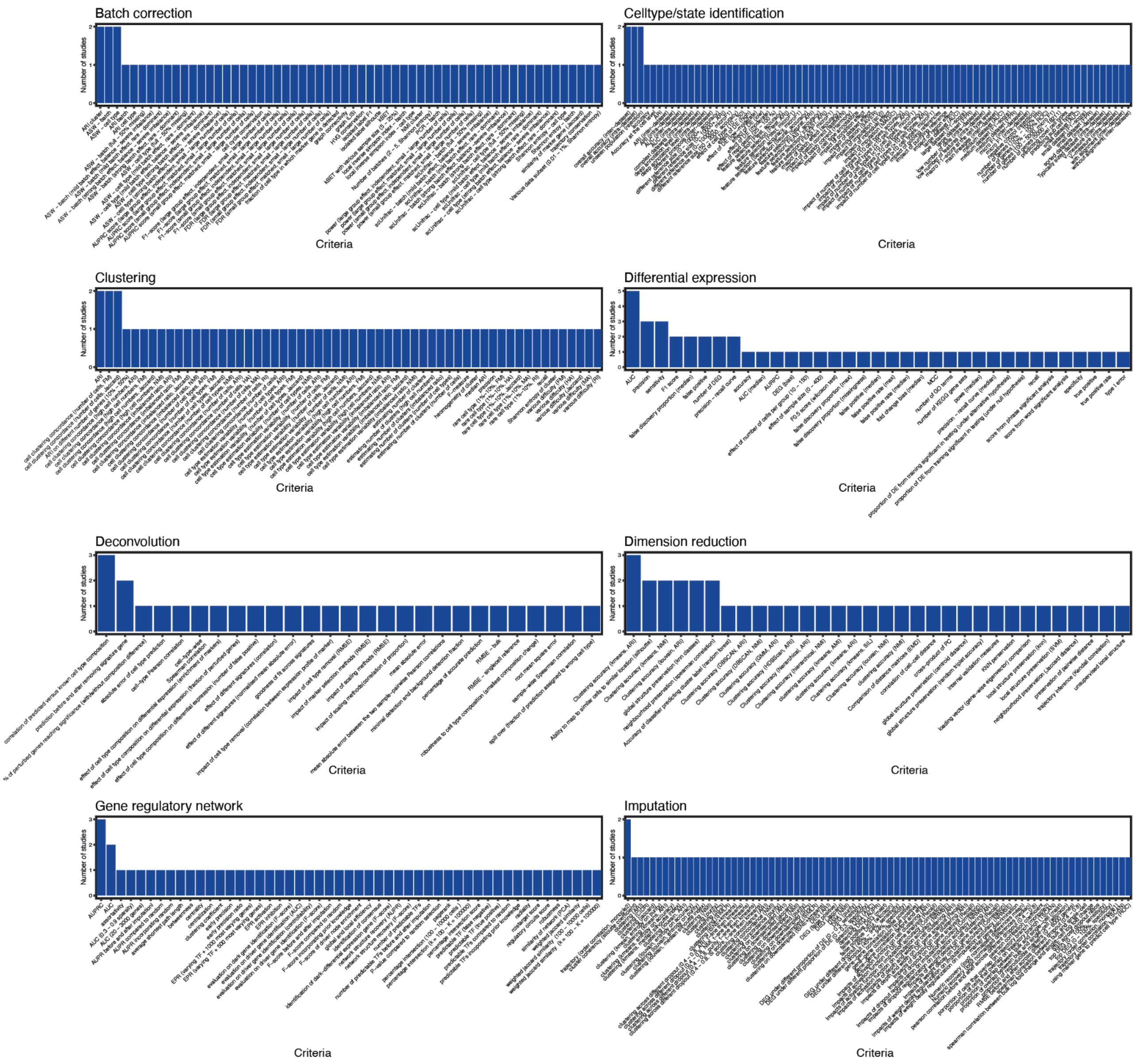


**Supplementary Figure 8. Overlap of the criteria used by multiple benchmarking studies.**

Topics with more than three benchmarking papers were assessed. The barplot denotes the number of papers that the criteria appeared in.

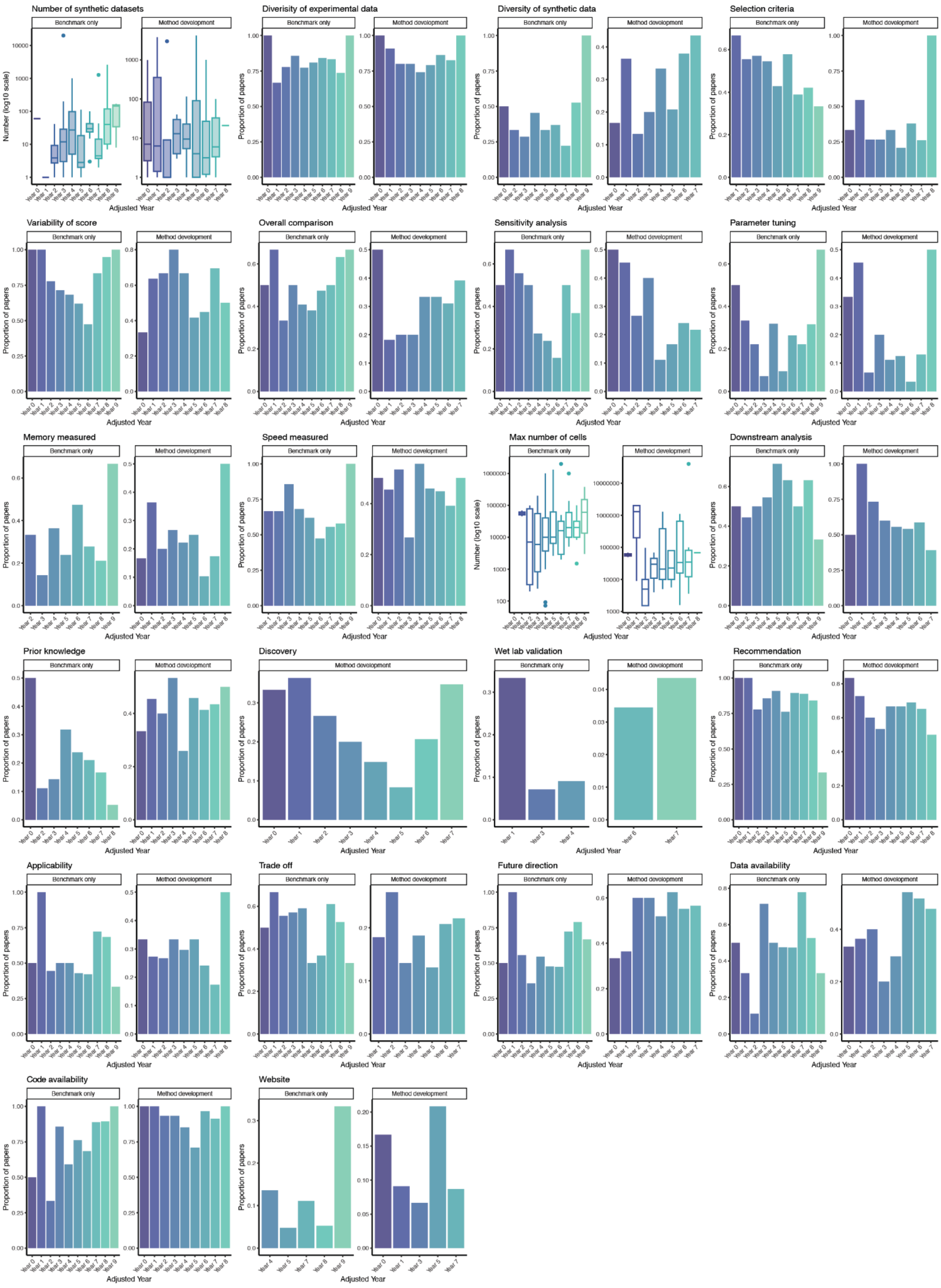


**Supplementary Figure 9. Trends of variables over adjusted year.**

This figure provides a visual overview of the evaluation criteria across the nine categories over the adjusted years.

### Supplementary Tables

**Supplementary Table 1: We would like to acknowledge the contribution of all the contributors listed below.**

| | **Contributor** |
|---|---|
| 1 | Adam Chan |
| 2 | Agus Salim |
| 3 | Andy Tran |
| 4 | Bárbara Zita Peters Couto |
| 5 | Beilei Bian |
| 6 | Carissa Chen |
| 7 | Chuhan Wang |
| 8 | Chunlei Liu |
| 9 | Daniel Kim |
| 10 | Daniel Mechtersheimer |
| 11 | Dantong Zhu |
| 12 | Dario Strbenac |
| 13 | Di Xiao |
| 14 | Farhan Ameen |
| 15 | Guan Gui |
| 16 | Hani Kim |
| 17 | Helen Fu |
| 18 | Ian Lee |
| 19 | Jean Yang |
| 20 | Lijia Yu |
| 21 | Marni Torkel |
| 22 | Nicholas Robertson |
| 23 | Sanghyun Kim |
| 24 | Shila Ghazanfar |
| 25 | Taiyun Kim |

| | |
|---|---|
| 26 | Terry Speed |
| 27 | Tian Lan |
| 28 | Wenze Ding |
| 29 | Xiaoqi Liang |
| 30 | Xiaoying Liao |
| 31 | Yingxin Lin |
| 32 | Yue Cao |
| 33 | Yunwei Zhang |

**Supplementary Table 2. Conversion of selected values into 0 to 1 for analytical purposes.**

| Details of variables | Original response | After conversion |
|---|---|---|
| All relevant variables involving “No”, “Yes” and “Not sure” | No | 0 |
| | Yes | 1 |
| | Not sure | 0.5 |
| Sensitivity | Default setting | 0 |
| | Parameter tuning | 1 |
| Types of dataset | Experimental data | 0.5 |
| | Synthetic data | 0.5 |
| | Both | 1 |
| Number of experimental datasets<br>Number of synthetic datasets<br>Number of methods compared<br>Max number of cells | Numeric value | Log10 normalisation, followed by scaling into [0, 1] |
